# Relationship Between Circulating FGF21 and Bone Mineral Density: An Exploratory Study

**DOI:** 10.1101/2025.02.16.638516

**Authors:** Matthew Peterson, Sydney Czanstkowski, Kathleen A. Richardson, LesLee Funderburk

## Abstract

**Introduction:** Fibroblast growth factor 21 (FGF21) has metabolic regulating effects on the body and exhibits endocrine-like effects on the body. Prior research has indicated that FGF21 may modulate bone density; however, the extent to which FGF21 can alter bone density and the effect of biological sex remains understudied.

**Methods:** A total of 25 individuals (Female = 13, Male =12) volunteered to have their whole body bone density measured using Dual X-Ray Absorptiometry (DEXA) and blood drawn on two separate occasions. Serum samples were analyzed for FGF21 concentration and concentration value was averaged between the two samples, all blood samples were drawn during the mid-follicular phase in female participants. The relationship between FGF21 and markers of bone density was examined using Pearson correlations.

**Results:** Whole-body BMD (wb-BMD) was negatively correlated with FGF21 across the whole sample (*r* = -0.53, *p* < 0.05). When stratifying the sample by biological sex, wb-BMD was negatively correlated with FGF21 in females (*r* = -0.72, *p* < 0.05) and in males (*r* = -0.18, *p* > 0.05); however, in males this relationship was not statistically significant.

**Conclusion:** FGF21 appears to be negatively correlated to bone density in humans with a significant relationship in females but not in males.

## Introduction

Fibroblast growth factor 21 (FGF21), has pluripotent, endocrine-like effects throughout the body. Most commonly known as a metabolic regulator, the effects of FGF21 on bone homeostasis remain understudied. Decreases in bone mineral density increase the fragility of bones along with the risk of fracture especially in aging individuals.(1) Previous research on the relationship between FGF21 and bone density is inconclusive.

In one study, ten obese rhesus macaque monkeys were administered increasing doses of pharmacological FGF21 for twelve weeks, with no adverse effects on bone density observed during the trial. (2) An observational study in humans used data from 115 healthy, postmenopausal women found no significant correlations between FGF21 and bone density.(3)

In contrast to the above, Lee et al. measured fasting plasma FGF21 levels and body composition among 40 volunteers.(4) Among the male volunteers, there was no significant correlation between FGF21 and bone density; however, in females, they observed a positive correlation between plasma FGF21 and bone density.(4) In mice, chronic exposure to FGF21 results in decreased bone mass.(5) Thus, the relationship between FGF21 and bone density remains incompletely understood.

Both bone density and FGF21 demonstrate sex specific effects.(6) Additionally, FGF21 has been shown to have a high intraday variability in humans,(7) which has the potential to confound investigations between FGF21 and bone density. Therefore, the purpose of this study was to investigate the relationship between FGF21 and bone density in healthy males and females who had blood FGF21 concentrations measured on two separate occasions.

## Methods

### Participants

As part of a larger investigation of FGF21 in humans, male and female participants were recruited for this study. Inclusion criteria included: age between 18 and 45 years old, regular exercisers (at least thirty minutes of exercise three days a week), a BMI between 18.5 and 24.9, and free from any cardiovascular or metabolic diseases. Additionally, female participants were eumenorrheic and not using hormonal contraceptives. All procedures were approved by the local institutional review board. All participants were informed of the study procedures and provided written informed consent prior to starting any study procedures.

### Bone Density

Bone density was measured using whole-body dual energy x-ray absorptiometry (DEXA) (Hologic Discovery Series W; Waltham, MA).

### FGF21

Due to the high inter-day variability in serum FGF21 concentrations,(7) blood was procured from each participant on two non-consecutive days following an overnight fast. In female participants, both blood draws were taken during the mid-follicular stage of the menstrual cycle (days 3-10).

Blood was allowed to sit at room temperature for thirty minutes and then centrifuged at 3,000 rpm for fifteen minutes. Serum samples were separated and stored at -80°C until analysis. Serum FGF21 concentrations were determined using commercially available enzyme-linked immunosorbent assays (ELISAs) (DF2100, R&D Systems). Average FGF21 concentration from the two blood samples was calculated prior to analysis.

### Statistics

A Shapiro-Wilk test was used to assess normality in all variables; FGF21 concentrations violated normality and were natural logarithm transformed prior to analysis. Pearson correlations were used to assess the relationship between FGF21 concentration and bone mineral density (BMD) and bone mineral content (BMC). All analyses were conducted using the R statistical software (R Foundation for Statistical Computing, Vienna, Austria). An alpha level of *p* < 0.05 was adopted throughout.

## Results

### Participant Characteristics

A total of twenty-five participants (female = 13, male = 12) were analyzed for this study. Demographic variables are provided in table 1.

**Table 1.** Demographics. BMI = body mass index. Data are presented as mean±SD.

| Variable | Whole Group | Females | Males |
| --- | --- | --- | --- |
| *n* | 25 | 13 | 12 |
| Height (m) | 1.7 ± 0.09 | 1.6 ± 0.06 | 1.8 ± 0.08 |
| Weight (kg) | 65.4 ± 11.07 | 57.9 ± 8.62 | 73.4 ± 7.06 |
| Age | 24.7 ± 5.56 | 22.6 ± 3.04 | 27.0 ± 6.81 |
| BMI | 22.3 ± 1.93 | 21.3 ± 1.92 | 23.4 ± 1.27 |

### FGF21 and Bone Density

Correlations were calculated between FGF21 and bone density variables. Statistically significant correlations for the whole sample were found between whole-body BMD (wb-BMD) (*r* = -0.53, *p* < 0.05), subtotal BMD (st-BMD) (*r* = -0.45, *p <* 0.05), BMD *z*-score (*r* = -0.48, *p* < 0.05), whole-body BMC (wb-BMC) (*r* = -0.45, *p* < 0.05), lumbar spine BMD (ls-BMD) (*r* = -0.62, *p* < 0.01), and pelvis BMD (p-BMD)(*r* = -0.57, *p* < 0.01)(table 2). Subtotal BMC (st-BMC) was not statistically significant for the whole group (*r* = -0.37, *p* > 0.05) (table 2).

**Table 2.** Whole Group Correlations. BMD = bone mineral density; subtotal BMD = BMD without the cranium; BMC = bone mineral content.

| Variables | *r* | *p*-value |
| --- | --- | --- |
| Subtotal BMD | -0.45 | <0.05 |
| Total BMD | -0.53 | <0.05 |
| BMD Z-Score | -0.48 | <0.05 |
| Subtotal BMC | -0.37 | 0.07 |
| Total BMC | -0.45 | <0.05 |
| Lumbar Spine BMD | -0.62 | <0.01 |
| Pelvis BMD | -0.57 | <0.01 |

When separating the sample population based on biological sex, all bone density variables were significantly and negatively correlated with blood FGF21 in females - wb-BMD (*r* = -0.72, *p* < 0.05), st-BMD (*r* = -0.71, *p* < 0.05), BMD *z*-score (*r* = -0.69, *p* < 0.05), wb-BMC (*r* = -0.70, *p* < 0.05), st-BMC (*r* = -0.67, *p* < 0.05), ls-BMD (*r* = -0.77, *p* < 0.01), p-BMD (*r* = -0.86, *p* < 0.01) (table 3). In males, there were no statistically significant correlations between blood FGF21 and bone density variables (see table 3).

**Table 3.** Sex Specific Correlations. BMD = bone mineral density; subtotal BMD = BMD without the cranium; BMC = bone mineral content.

| Variable | Female *r* | Female *p*-value | Male *r* | Male *p*-value |
| --- | --- | --- | --- | --- |
| Subtotal BMD | -0.71 | <0.05 | 0.03 | 0.92 |
| Total BMD | -0.72 | <0.05 | -0.18 | 0.57 |
| BMD Z-Score | -0.69 | <0.05 | -0.16 | 0.61 |
| Subtotal BMC | -0.67 | <0.05 | 0.17 | 0.60 |
| Total BMC | -0.70 | <0.05 | -0.01 | 0.97 |
| Lumbar Spine BMD | -0.77 | <0.01 | -0.41 | 0.19 |
| Pelvis BMD | -0.86 | <0.01 | 0.05 | 0.88 |

## Discussion

In the present study, we observed a negative correlation between measurements of bone health and FGF21 in young, apparently healthy humans. When analyzing the results by biological sex, a negative relationship between FGF21 and markers of bone health was found exclusively in females, but not in males. This suggests that FGF21 may be related to bone health in females, but not in males. The conclusions of this study add to the literature that FGF21 exhibits sex-specific effects in humans.(6)

Prior research into the relationship between FGF21 and bone density in animal models has been mixed. In one study, ten obese (five males and five females), non-diabetic, rhesus macaque monkeys were administered increasing doses of pharmacological FGF21 while maintaining a high fat diet for twelve weeks.(2) The results showed no adverse effects on bone density over the course of the trial, but did see a 2-fold increase in a marker of bone resorption known as CTX-1.(2) However, Wei et al. examined mice and the consequences of chronic FGF21 exposure on bone.(5) They measured bone markers by ELISA and found that the bone formation marker N-terminal propeptide was 40% lower and the bone resorption marker C-terminal telopeptide was 122% higher in the FGF21-transgenic mice indicating that FGF21 does influence bone homeostasis and decreases bone mass.(5)

Similarly mixed results have been found in humans. One study measured fasting plasma FGF21 concentrations and BMD finding that there was a positive correlation between FGF21 and BMD in females, but not in males.(4) In contrast, a different study with a mixed sample of males and females found a negative relationship between blood FGF21 and BMD.(8) Hao et al., reported a negative relationship between bone density and circulating FGF21, but did not find any sex specific differences.(9) Finally, a study performed by Choi et al. in post-menopausal women found no relationship between BMD and FGF21 concentrations.(3) Thus, a variety of relationships between FGF21 and humans have been reported. The present study adds to the current body of literature by reporting that there is a negative relationship between FGF21 and BMD in healthy humans, and that this relationship is sex-specific.

One confounding factor when investigating FGF21 in humans is that blood FGF21 concentrations exhibit a high degree of day to day variability.(7,10) Thus measurements of FGF21 concentrations on a particular day may not accurately reflect the average concentrations present in a person. One advantage of the current study over prior studies is that we measured FGF21 on two separate occasions, increasing the likelihood that our measurements accurately captured the typical FGF21 concentrations found in our participants. However, the current study is not without limitations. First, bone density was measured using a whole-body DEXA scan, rather than the segmental scans that are used clinically to diagnose osteoporosis. Additionally, we relied upon a small sample of young, apparently healthy individuals; therefore, the results might not translate to other populations.

## Conclusions

The primary results of this present study are that there is a negative relationship between FGF21 and bone density in healthy humans and this relationship is sex specific. Future studies in the area of FGF21 and bone density could investigate the relationship longitudinally to determine if this relationship remains consistent across time. Additionally, randomized trials with exogenous FGF21 administration would also be beneficial for determining a cause-and-effect relationship.

